# Peptide-MHC-targeted retroviruses enable *in vivo* expansion and gene delivery to tumor-specific T cells

**DOI:** 10.1101/2024.09.18.613594

**Authors:** Ellen J.K. Xu, Blake E. Smith, Winiffer D. Conce Alberto, Michael J. Walsh, Birkley Lim, Megan T. Hoffman, Li Qiang, Jiayi Dong, Andrea Garmilla, Qingyang Henry Zhao, Caleb R. Perez, Stephanie A. Gaglione, Connor S. Dobson, Michael Dougan, Stephanie K. Dougan, Michael E. Birnbaum

## Abstract

Tumor-infiltrating-lymphocyte (TIL) therapy has demonstrated that endogenous T cells can be harnessed to initiate an effective anti-tumor response. Despite clinical promise, current TIL production protocols involve weeks-long *ex vivo* expansions which can affect treatment efficacy. Therefore, additional tools are needed to engineer endogenous tumor-specific T cells to have increased potency while mitigating challenges of manufacturing. Here, we present a strategy for pseudotyping retroviral vectors with peptide-major histocompatibility complexes (pMHC) for antigen-specific gene delivery to CD8 T cells and examine the efficacy of these transduced cells in immunocompetent mouse models. We demonstrate that pMHC-targeted viruses are able to specifically deliver function-enhancing cargoes while simultaneously activating and expanding anti-tumor T cells. The specificity of these viral vectors enables *in vivo* engineering of tumor-specific T cells, circumventing *ex vivo* manufacturing processes and improving overall survival in B16F10-bearing mice. Altogether, we have established that pMHC-targeted viruses are efficient vectors for reprogramming and expanding tumor-specific populations of T cells directly *in vivo*, with the potential to substantially streamline engineered cell therapy production for a variety of applications.

## INTRODUCTION

Adoptive cell therapy is a promising cancer treatment modality that uses expanded or genetically modified T cells to recognize and potently kill tumor cells. Tumor infiltrating lymphocyte (TIL) therapy was approved by the FDA in 2024 for advanced melanoma, becoming the first cell therapy approved for solid tumors (*1*). TIL therapies are produced by isolating autologous T cells from tumor biopsies and extensively expanding the cells *ex vivo* before reinfusing them into the patient. Early exploration of TIL therapy highlighted its ability to induce long-lasting remission in patients with metastatic melanoma (*2, 3*), and since then, varying frequencies of TILs have been detected in a range of cancers (*4*–*7*), underscoring the potential benefit of using these cells as a treatment for multiple cancer types.

The phenotype and quality of TIL products have been demonstrated to significantly influence clinical outcomes. Prior to reinfusion, TILs often exhibit exhausted phenotypes due to their previous intratumoral antigen exposure and the prolonged *ex vivo* expansion protocol, impeding them from mounting an aggressive and durable anti-tumor response (*8*). To mitigate this, additional engineering strategies to improve cell potency could be incorporated into production protocols with the goals of delivering immunostimulatory cargoes (*9, 10*) and diminishing immunosuppressive signals (*11*–*13*). However, current methods to introduce these modifications would apply them to all T cells present in the TIL expansion, increasing potential toxicities resulting from engineered bystander T cells while further increasing manufacturing complexity. Evidence from other engineered cell therapies suggests that the persistence and efficacy of adoptively transferred cells diminishes as the interval between T cell isolation and reinfusion increases (*14, 15*), reinforcing the need to minimize production time in order to increase treatment success. Additional strategies are therefore desirable to revitalize and generate a more effective anti-tumor response before engineered TIL therapies can be widely translated to the clinic.

Developing gene delivery vectors to modify tumor-specific T cells directly *in vivo* could mitigate many of these challenges involving *ex vivo* TIL manufacturing and potency. Ongoing efforts to generate viral vectors for *in vivo* gene delivery are focused on limiting viral tropism to specific T cell lineage markers such as CD3, CD8, CD4, CD62L, and CD5 (*16*–*23*). While these strategies may be appropriate for the generation of chimeric antigen receptor (CAR) T cells, since they will modify broad subsets of T cells, they are less appropriate for antigen-specific transduction which could enhance the efficacy of TIL therapies. Recently, targeted gene delivery to antigen-specific T cells has been demonstrated, relying on display of peptide-major histocompatibility complex (pMHC) to be recognized by a T cell clone’s unique T cell receptor (TCR) (*24*–*26*). The development of lipid nanoparticles targeted by pMHC have demonstrated that *in vivo* antigen-specific delivery of mRNA-encoded cargoes is possible (*25*), yet longer term transgene expression will likely be required to ensure robust anti-tumor responses. While there is some precedent that functional cargoes can be delivered using lentiviral vectors via TCR-pMHC interactions *in vitro* (*26*), no studies have evaluated the efficacy of viruses displaying pMHC (pMHC-targeted viruses) as anti-tumor immunotherapeutic agents.

Here, we demonstrate that pMHC-targeted viruses can be used to efficiently reprogram anti-tumor T cell populations, delivering a therapeutic cargo that generates an effective anti-tumor response *in vivo*. First, we developed a pMHC-targeted gammaretrovirus pseudotyping strategy to facilitate efficient gene delivery to murine antigen-specific T cells in polyclonal populations, enabling evaluation of treatment efficacy in immune competent models. We demonstrated our approach using a tethered version of the cytokine interleukin-12 (IL-12) as a test cargo, given its potent immunomodulatory effects but dose-limiting toxicities when delivered systemically (*27, 28*). Transducing antigen-specific T cells *in vitro*, we observed activation and expansion of target T cells that were able to extend survival in a B16F10 melanoma model. Finally, we determined that pMHC-targeted viruses delivering tethered IL-12 could be infused directly *in vivo* to generate long-lived, memory T cells that extend tumor control. Taken together, we demonstrate that pMHC-targeted viruses are safe and effective vehicles for *in vivo* gene delivery to anti-tumor T cells, with the potential to customize these vectors for a wide range of antigen-specific T cell modifications.

## RESULTS

### Peptide-MHC pseudotyped viruses efficiently activate, expand, and transduce murine CD8 T cells

Our lab and others have demonstrated that lentiviruses can be pseudotyped via co-expression of pMHC and a receptor-blinded version of the vesicular stomatitis glycoprotein (termed “VSVGmut”) that can serve as a fusogen to enable antigen-specific transduction of human T cells (*24, 26, 29*). To evaluate the potential of pMHC-targeted viruses as gene delivery vectors, we first established a murine model to study the full scope of function of virally engineered antigen-specific T cells. Rather than rely on xenograft models, which would restrict evaluation of cargoes to those that could function without a complete immune system, we chose to adapt pMHC-targeted vectors for use in syngeneic models.

To demonstrate our approach, we used TRP1^high^ TCR transnuclear mice as a source of CD8 T cells that recognize a well-described melanoma antigen, tyrosinase-related protein 1 (TRP1), with physiological pMHC:TCR affinity (*30*). Based on previous studies using the TRP1^high^ system, we selected the strong agonist peptide mimotope of the TRP1 epitope, AAPDLGYM, presented by the murine class I MHC H2-D^b^ in the format of a single chain pMHC trimer (termed “A1”), as an initial targeting molecule for proof-of-concept experiments (*31*–*33*). Given previously reported limitations of using lentiviruses to efficiently infect murine T cells (*34*), we adapted our targeting approach for use with murine leukemia gammaretrovirus (MuLV) as the viral vector for co-display of pMHC and fusogen (Fig.1A).

We hypothesized that pMHC-displaying viral particles would selectively activate antigen-specific target cells in a population of naïve T cells. Therefore, we evaluated the efficiency of A1-targeted viruses displaying unmutated VSVG, VSVGmut, or the ecotropic envelope protein (Eco) derived from the Moloney murine leukemia virus, a commonly implemented gammaretroviral vector for gene delivery to pre-activated murine CD8 T cells (*35*–*38*). Targeted gammaretroviruses delivering a fluorescent cargo (ZsGreen), each displaying A1 and a fusogen, were used to transduce mixtures of freshly isolated on-target (TRP1^high^) and off-target, polyclonal wild-type C57BL/6J T cells (B6) murine CD8 T cells. After two days, the highest on-target transduction was mediated by viruses co-expressing A1 with the ecotropic envelope fusogen (A1-ZsG) (Fig. 1B). Viruses targeted with a non-cognate pMHC (displaying an ovalbumin-derived peptide) or Eco alone did not mediate transduction in unactivated cells (Fig. 1C). However, when added to pre-activated cells, the untargeted virus displaying only Eco (Eco-ZsG) was able to induce robust transduction of both TRP1^high^ and B6 CD8 T cells, validating that this parental virus was functional and not antigen-specific (fig. S1).

**Fig. 1.**
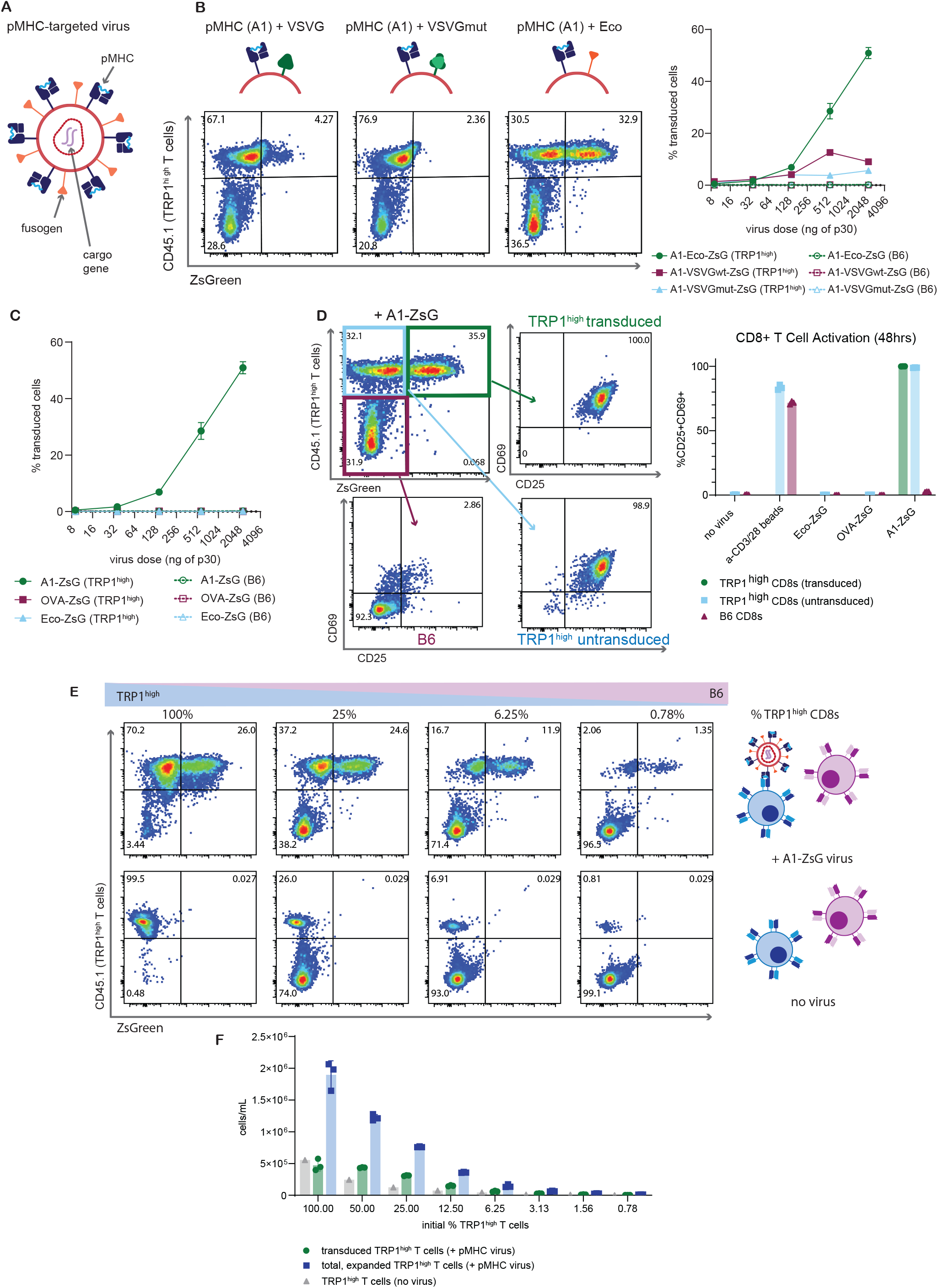
Viral vectors co-expressing pMHC and the MuLV ecotropic envelope selectively activate, expand, and transduce antigen-specific murine CD8 T cells. (**A**) Schematic of pMHC-targeted gammaretrovirus detailing the co-expression of the targeting (pMHC) and fusogen molecules on the viral membrane. The therapeutic cargo to be delivered is packaged in the viral genome. (**B**) Different pMHC + fusogen combinations were tested to determine the most efficient pseudotype (depicted by each of the cartoons). ZsGreen-expressing viruses were added to a mixture of 50% on-target (TRP1^high^) and 50% off-target (B6) CD8s with 8ug/mL polybrene and spinfected. Transduction was assayed after 48 hours, with representative flow plots shown. (**C**) pMHC-targeted viruses displaying an on-target pMHC (A1-ZsG), off-target pMHC (OVA-ZsG), or no pMHC (Eco-ZsG) were added to a mixture of 50% on-target (TRP1^high^) and 50% off-target (B6) CD8s with 8ug/mL polybrene and spinfected. Percent ZsGreen+ cells assessed via flow cytometry 48 hours after addition of virus. (**D**) Representative plot from (B) showing expression of activation markers CD25 and CD69 in TRP1^high^ transduced, TRP1^high^ untransduced, and off-target B6 CD8s measured 48 hours after addition of A1-Eco-ZsGreen virus (2048ng p30 dose). Summary plot showing percent CD25+CD69+ after addition of different pMHC-targeted viruses or anti-CD3/CD28 beads. (**E**) Representative flow plots depicting different starting frequencies of TRP1^high^ T cells mixed with B6 CD8s (bottom row) and the resultant expansion and transduction (top row) after transducing those mixtures with A1-ZsG virus. The same dose (924ng p30) of A1-ZsG virus was added to all populations. Transduction efficiency was assessed after 48 hours via flow cytometry. (**F**) Absolute cell counts from experiment in (E). Grey bars represent conditions with no virus added. Green bars indicate absolute counts of transduced TRP1^high^ T cells after dosing cell mixtures with A1-ZsG virus. Blue bars quantify expansion of total, antigen-specific TRP1^high^ population when A1-ZsG virus was added.

We observed specific activation of on-target TRP1^high^ T cells when A1-targeted viruses were added to the CD8 T cell mixture, whereas off-target cells in the same well were not activated (Fig. 1D). Addition of untargeted Eco-ZsG or an off-target pMHC-ZsG virus did not result in similar levels of CD69 or CD25 expression in TRP1^high^ CD8 T cells compared to addition of A1-ZsG. Even in TRP1^high^ T cells that were not transduced to express ZsGreen, we observed similar proportions of CD25+CD69+ CD8 T cells compared to transduced TRP1^high^ T cells, indicating that pMHC-displaying viruses are potent activators of anti-tumor T cells.

We next evaluated the efficiency of A1-ZsG viruses at selectively transducing on-target cells at lower initial frequencies. In patient samples, the endogenous frequency of anti-tumor, cytotoxic T lymphocytes ranges from ∼1% for MART-1 specific T cells to less than 0.01% for WT-1, MAGE family, and NYESO-1 specific T cells (*39*). We diluted TRP1^high^ CD8 T cells into B6 CD8s at different ratios and found that A1-ZsG viruses were able to transduce on-target cells at all starting frequencies (Fig. 1E). Our pMHC-targeted viruses were able to expand rare, cognate T cells (Fig. 1F), suggesting that pMHC-targeted viruses could be used to deliver therapeutic cargoes to enhance anti-tumor T cell function with the additional benefit of simultaneously expanding the overall number of anti-tumor, effector T cells.

To validate the generality of our pMHC targeting approach, we next demonstrated that pMHC-targeted viral vectors can be readily adapted for other antigen targets. To this end, we generated single-chain trimer versions of H2-K^b^ presenting the ovalbumin-derived model antigen SIINFEKL (OVA) (*40*), and H2-L^d^ presenting QLSPFPFDL (QL9), an agonist for the 2C TCR model system (*41, 42*). Pseudotyping viruses with each of these pMHC plus Eco, we observed that the vectors maintained high specificity for their cognate TCR compared to off-target B6 CD8s (Fig. 2, A and B). Furthermore, in each of these additional systems, pMHC-targeted viruses both expanded and transduced on-target T cells (Fig. 2, C and D). Taken together, we have developed a modular pMHC-based pseudotyping strategy that enables specific activation, expansion, and transduction of murine antigen-specific T cells from rare, polyclonal populations *in vitro*.

**Fig. 2.**
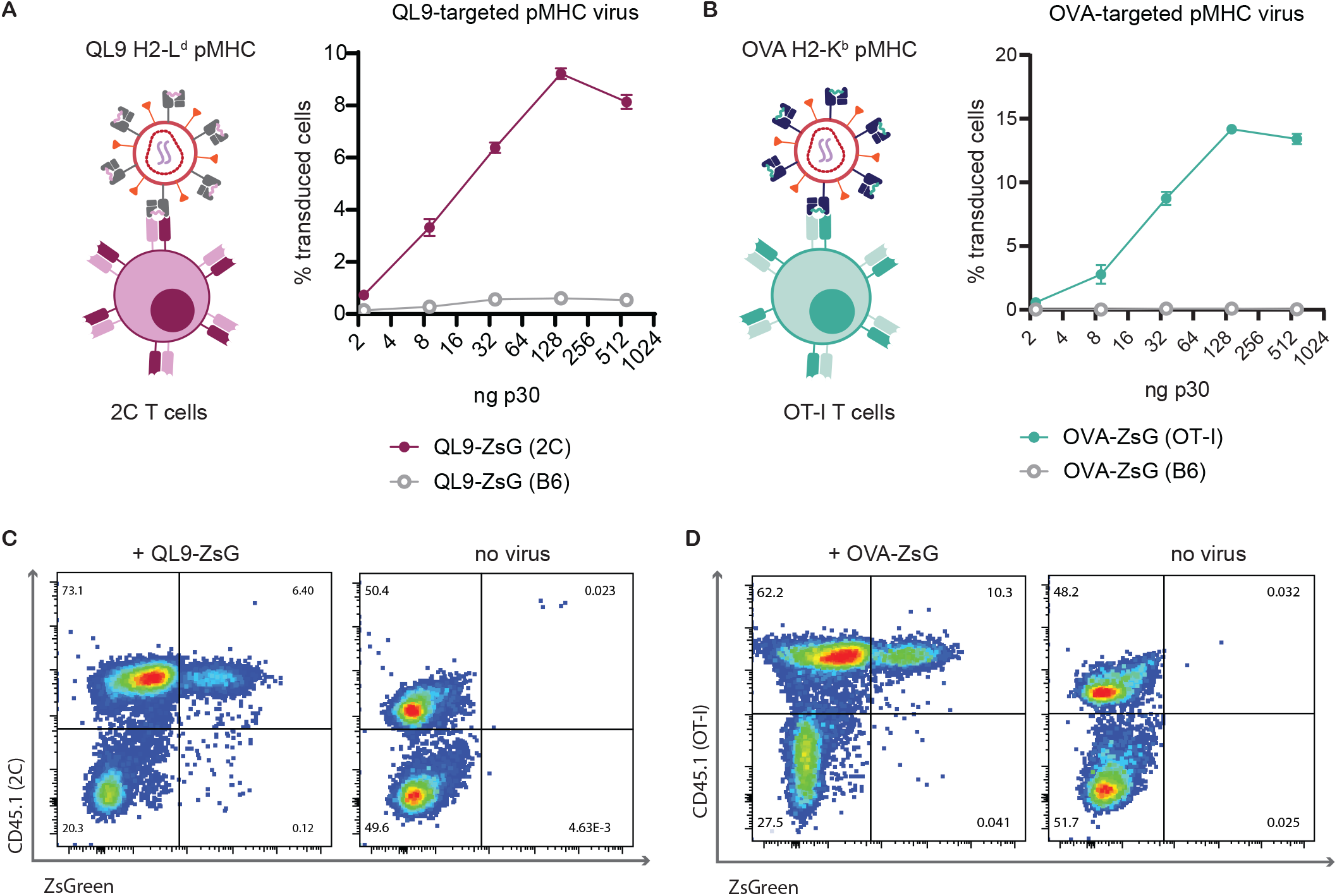
pMHC-targeted viruses are versatile tools for gene delivery to antigen-specific T cells in a variety of model systems. (**A**) + (**B**) pMHC-targeted gammaretroviruses can be retargeted to different antigen-specific T cell populations by changing the single chain trimer pMHC that is co-expressed with the ecotropic envelope glycoprotein. Two different ZsGreen viruses were generated, displaying a QL9 or OVA peptide in different MHC alleles to target 2C or OT-1 T cells respectively. Percent transduction as measured after 48 hours post virus addition is shown for QL9-ZsG and OVA-ZsG viruses added to a 1:1 ratio of on- and off-target cells, where cognate transgenic TCR T cells were used as the on-target population and B6 CD8s were used as the off-target population. Data representative of two separate experiments. (**C**) + (**D**) Representative flow plots from (A) + (B) where on-target virus (dosed at 512ng p30) was added to a 1:1 mixture of on- and off-target cells (left) compared to a control well (right) where no virus was added.

### A1-targeted viruses can deliver a functional, tethered mIL-12 cargo

Many gene cargoes have been previously reported to enhance the potency, trafficking, or safety of adoptively transferred engineered T cell therapies (*43*). We hypothesized that our pMHC-targeted viruses would be particularly useful for delivery of cargoes reported to have robust anti-tumor effects that are limited by systemic toxicities. We selected a tethered murine interleukin-12 (mIL-12) construct as a proof-of-concept cargo for genetic manipulation of antigen-specific T cells whose efficacy had previously been validated in conjunction with CAR T cell therapy (*44*) (Fig. 3A). We confirmed that IL-12 is detectable on the surface of cells transduced with targeted A1-mIL12 viruses using an anti-murine IL-12 antibody (Fig. 3B). When A1 viruses encoding mIL-12 were added to a mixed population of on- and off-target CD8 T cells, A1-targeted vectors maintained specificity for TRP1^high^ T cells as demonstrated by selective expansion of on-target cells (fig. S2). IL-12 signaling drives interferon gamma (IFNγ) secretion, which was detectable in the supernatants of all conditions transduced with IL-12 viruses (Fig. 3, C and D). These results further highlight the specificity of A1-targeted viruses, as IFNγ secretion is significantly elevated when A1-mIL12 virus was added to TRP1^high^ T cells but not to polyclonal B6 T cells (Fig. 3C). In contrast, untargeted Eco-mIL12 virus transduced and elicited IFNγ secretion in both TRP1^high^ and B6 T cells (Fig. 3D). Increased pSTAT4 signaling was also observed in TRP1^high^ cells transduced with A1-mIL12 virus compared to A1-ZsG virus, demonstrating that the mIL-12 construct initiates expected downstream signaling cascades in transduced T cells (Fig. 3E). Interestingly, elevated pSTAT4 levels were also detected in untransduced TRP1^high^ T cells, indicating that the tethered cytokine is able to mediate effects *in trans*. Taken together, these data demonstrate that pMHC-targeted viruses are able to specifically deliver a functional cargo *in vitro*.

**Fig. 3.**
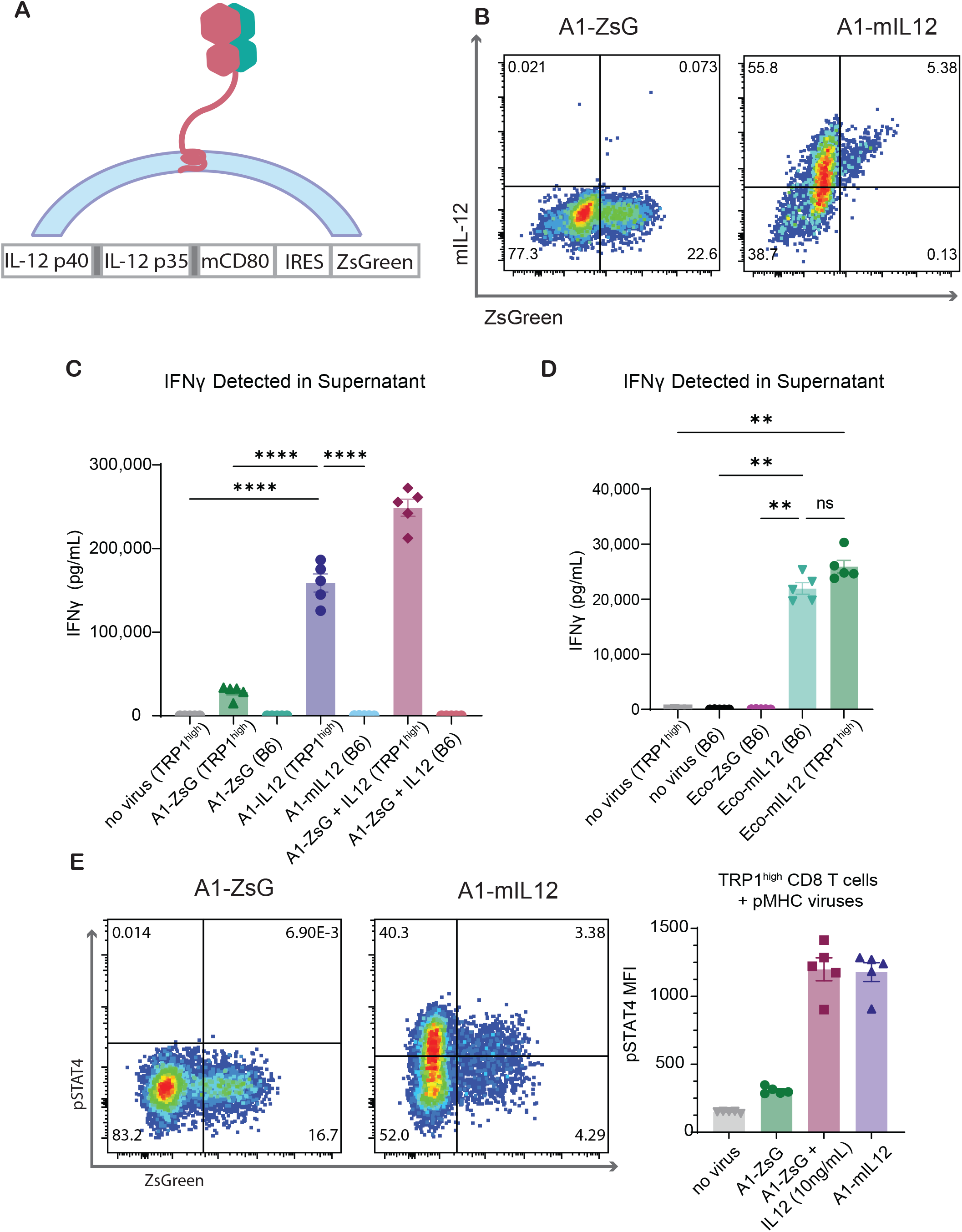
pMHC-targeted retroviruses enable delivery of a functional cargo, tethered IL-12, to tumor-specific T cells. (**A**) Schematic and construct design of tethered IL-12. The shaded grey boxes represent flexible linkers. (**B**) Representative flow plots of transduction achieved with A1-ZsG or A1-mIL12 viruses added to TRP1^high^ CD8 T cells after 48 hours. Single experiment from *n* = 3 biological replicates. (**C**) IFNγ secretion in supernatants from *in vitro* cultures of freshly isolated TRP1^high^ CD8 T cells transduced with A1-targeted viruses was quantified via ELISA. Supernatants were collected 48 hours after transduction. Virus doses were standardized to 1.94E+09 particles per 150,000 cells. In the A1-ZsG + IL12 condition, cells were resuspended in media supplemented with 10ng/mL murine IL-12 and transduced with A1-ZsG. P values were calculated by Bonferroni-corrected one-way ANOVA. ****p < 0.0001 (**D**) Results from an IFNγ ELISA of supernatants from *in vitro* cultures of pre-activated cells transduced with Eco-targeted viruses. Supernatants were collected 48 hours after transduction. Virus doses were matched to those used in (C). Significance was determined using Bonferroni-corrected one-way ANOVA. **p < 0.01 (**E**) TRP1^high^ CD8 T cells were isolated and transduced with A1-ZsG or A1-mIL12 virus. 48 hours after transduction, cells were fixed, permeabilized, and stained with anti-pSTAT4 antibody before subsequent analysis via flow cytometry. Cells were transduced with A1-ZsG virus and cultured in media with 10ng/mL IL-12 as a positive control.

### Anti-tumor T cells transduced with tethered IL-12 *ex vivo* extend survival in B16F10 tumor-bearing mice

Having demonstrated the ability of pMHC-targeted viruses to deliver immunostimulatory cargoes *in vitro*, we explored how these transduced cells function in a therapeutic context. We selected B16F10 melanoma as an aggressive tumor model system that endogenously expresses the TRP1 antigen (*45, 46*). After inoculating B16F10 tumor cells subcutaneously in B6 mice, we harvested donor CD8 T cells from transgenic TRP1^high^ mice and transduced the cells *ex vivo* with either A1-ZsG, A1-mIL12, or untargeted Eco-ZsG (Fig. 4A). As an additional control, we transduced activated B6 CD8s with Eco-mIL12 virus to determine whether delivery of IL-12 to polyclonal T cells would be sufficient for therapeutic efficacy. After two days *in vitro*, the cell product from each group was assessed for baseline transduction and activation state prior to adoptive transfer (fig. S3).

**Fig. 4.**
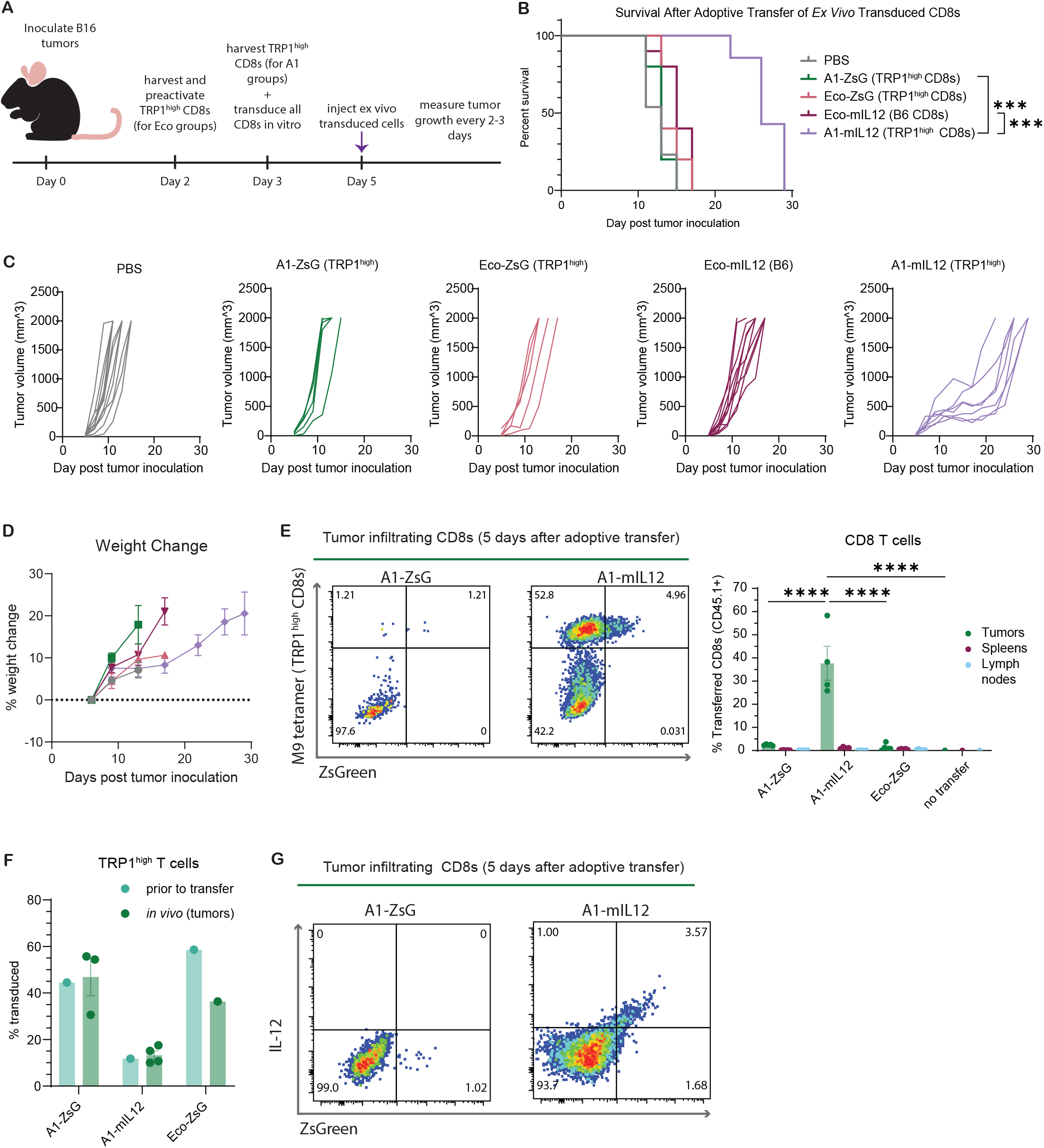
Adoptive transfer of TRP1^high^ T cells transduced by pMHC-targeted viruses delivering tethered IL-12 extends survival of B16F10 tumor-bearing mice. (**A**) 500,000 B16F10 tumor cells were inoculated on the dorsal flank of B6 mice. Donor TRP1^high^ CD8 T cells were isolated and transduced *ex vivo* with targeted and non-targeted virus. Two days later, percent transduction was quantified for each group and cells were transferred to recipient mice. Mice were subsequently monitored for tumor growth and signs of morbidity. (**B**) Kaplan-Meier curve for overall survival. Results from one experiment where *n* = 3 biological replicates. P values were determined using a Bonferroni-corrected log-rank (Mantel-Cox) test. *** p < 0.001 (**C**) Individual growth curves from mice in (B). Tumor size was measured in three dimensions (length, width, height) every 2-4 days and plotted as tumor volume. (**D**) Mouse weights were monitored every 2-4 days and change in weight was normalized to one day after cell injection. (**E**) Representative plots of tumor-infiltrating CD8 T cells harvested at 5 days after adoptive transfer of *ex vivo* transduced cells, gated on live CD8s. Summary plot of transferred cells detected in each of the three tissues harvested within each group includes mean ± SEM. P values were calculated by Bonferroni-corrected two-way ANOVA. **** p < 0.0001 (**F**) Quantification of transduced cells as a percentage of total TRP1^high^ T cells, comparing the percent transduced detected *in vivo* to initial percent transduced *ex vivo* prior to infusion. Any samples where number of transferred cells detected was less than 10 were excluded from analysis. Mean ± SEM plotted. (**G**) Representative panels of IL-12 and ZsGreen expression in tumor-infiltrating CD8 T cells gated on live CD8s for samples from (E). Each plot represents one mouse.

Consistent with previous reports, TRP1^high^ T cells alone were not sufficient to control B16F10 tumors (*30, 31, 33, 47*). However, mice that received TRP1^high^ T cells transduced with A1-mIL12 virus were able to maintain smaller tumors at early timepoints, ultimately experiencing a significant survival benefit (Fig. 4, B and C). In contrast, delivery of tethered IL-12 to polyclonal CD8s in the Eco-mIL12 group did not lead to the same prolonged survival, indicating that tethered IL-12 must be specifically delivered to anti-tumor T cells for greatest therapeutic efficacy. While previous work has demonstrated that injections of soluble IL-12, even if given intratumorally, causes drastic initial weight loss in mice(*48*), no weight loss or other evidence of toxicity was observed in mice treated with TRP1^high^ T cells transduced with tethered mIL-12 via A1-targeted viruses (Fig. 4D).

Given the improvement in overall survival mediated by the transfer of A1-mIL12 transduced cells, we next validated whether transduced cells could traffic to and persist in tumors. Following the same adoptive transfer protocol, TRP1^high^ T cells were transduced *ex vivo* and transferred into B16F10 tumor-bearing mice. Five days after transfer, tumors, spleens, and tumor-draining lymph nodes were harvested for analysis of infiltrating transferred and transduced cells. Across the tissues sampled, we detected an increased percentage of adoptively transferred CD8 T cells in tumors from the A1-mIL12 transduced group compared to other experimental groups and in comparison to other tissues within the A1-mIL12 treatment group (Fig. 4E). These results indicate that TRP1^high^ T cells are able to preferentially and efficiently traffic to tumors. In all virus transduced groups, the fraction of transferred cells detected that were transduced (ZsGreen+) mirrored the baseline frequency of ZsGreen+ cells prior to adoptive transfer (Fig. 4F), verifying that gene delivery via pMHC-targeted viruses does not limit *in vivo* persistence of transduced cells relative to untransduced cells. Additionally, tethered mIL-12 expression on tumor-infiltrating, transduced CD8s was confirmed in the A1-mIL12 transduced group via flow cytometry (Fig. 4G). Taken together, these results demonstrate that A1-mIL12 transduced cells are able to survive and traffic to tumors *in vivo* where they mediate prolonged survival in an aggressive, syngeneic tumor model.

### Intravenously delivered pMHC virus targets naïve CD8 T cells *in vivo*

Considering the specificity of A1-targeted viruses in polyclonal populations *in vitro*, we investigated whether pMHC-targeted viruses could target anti-tumor T cells directly *in vivo*. B6 mice partially reconstituted with TRP1^high^ T cells were treated with A1- or Eco-targeted virus intravenously (i.v.) (Fig. 5A). A1-mIL12 monotherapy mice or those receiving A1-mIL12 in combination with anti-PD1 antibody (aPD1) experienced a significant extension in overall survival compared to mice treated with A1-ZsGreen or Eco-mIL12 virus, underscoring that specific delivery of the therapeutic cargo to tumor-specific T cells was critical for therapeutic benefit (Fig. 5, B and C). Despite a modest increase in survival for A1-mIL12 + aPD1 compared to A1-mIL12, we observed no significant difference between these groups, suggesting that PD-1 was not primarily responsible for the eventual loss of immune-mediated control.

**Fig. 5.**
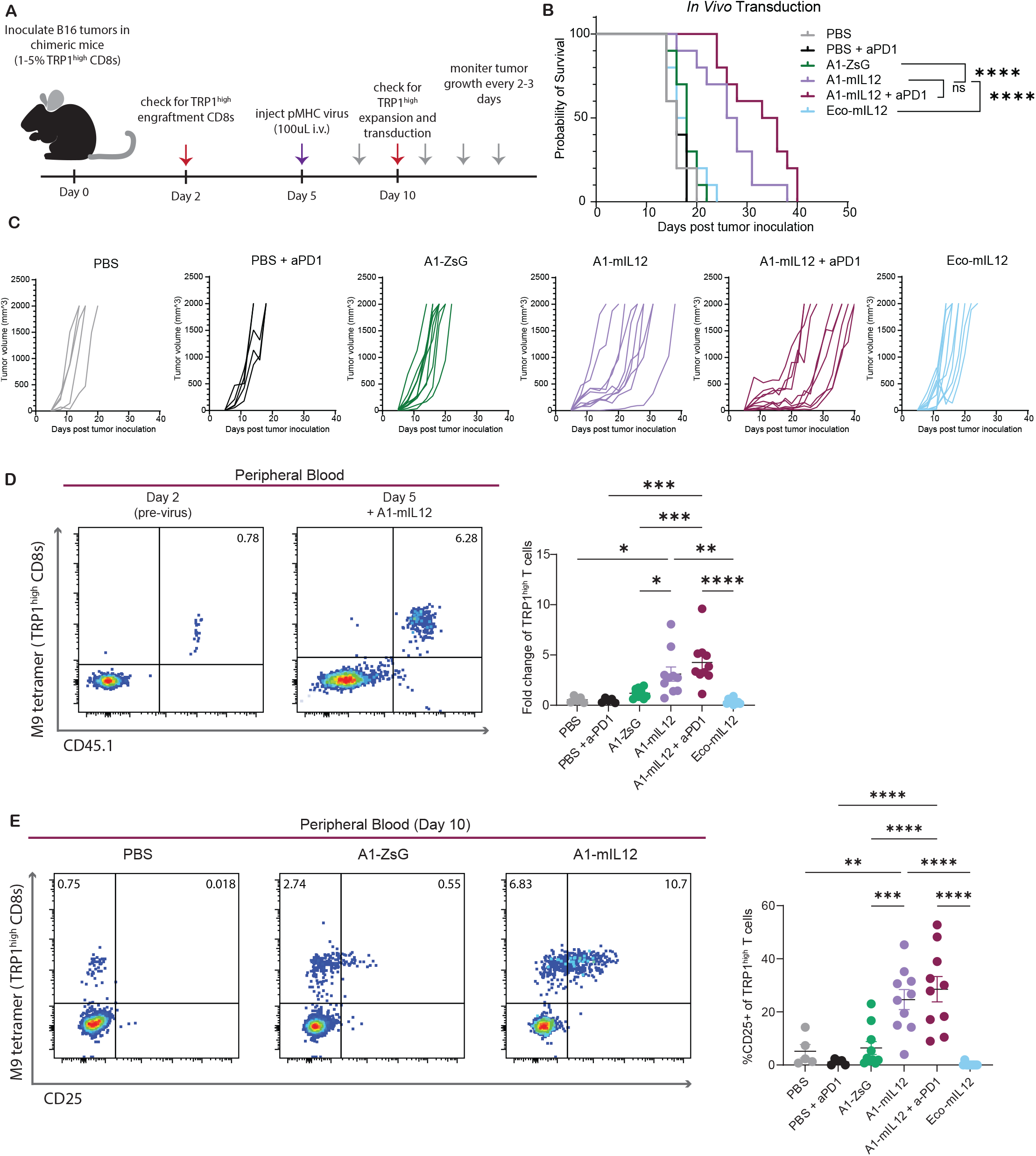
pMHC-targeted viruses delivering mIL12 generate *in vivo* engineered anti-tumor T cells and extend survival in a syngeneic, solid tumor model. (**A**) B6 mice were irradiated and partially reconstituted with TRP1^high^ lymphocytes. Two days later, 500,000 B16F10 tumor cells were inoculated subcutaneously in the dorsal flank. Three days later, 100uL of targeted viruses were injected i.v. via the tail vein. Grey arrows indicate i.p. injections of anti-PD-1 antibody (150ug) on days 8, 11, 14, and 17. Red arrows indicate blood draws to evaluate TRP1^high^ T cell engraftment prior to virus injection as well as expansion/transduction post-virus injection. (**B**) Kaplan-Meier curve for overall survival. Representative of two biological replicates where *n* = 10 mice per group. P values were determined by a Bonferroni-corrected log-rank (Mantel-Cox) test. ****p < 0.0001 (**C**) Tumor size was measured in three dimensions (length, width, height) every 2-4 days and plotted as tumor volume for individual mice. (**D**) Expansion of TRP1^high^ T cells calculated based on the percentage of tetramer+ and CD45.1+ cells in the peripheral blood, where CD45.1 is a marker for transferred TRP1^high^ lymphocytes. Each flow cytometry plot is representative of one mouse and is gated on live CD8 T cells. Annotated percentages are of total, live CD8 T cells. P values were calculated by Bonferroni-corrected one-way ANOVA. *p < 0.05, ***p < 0.001 (**E**) Expression of CD25 in peripheral blood samples from Day 10, gated on live CD8s. Each plot is representative of one mouse, and annotated percentages are of total, live CD8 T cells. P values were calculated via Bonferroni-corrected one-way ANOVA. ** p < 0.01, **** p < 0.0001

Comparing TRP1^high^ T cell frequencies in the peripheral blood before and after virus injection, we saw the greatest fold expansion of TRP1^high^ T cells following injection of A1-mIL12 virus compared to A1-ZsG, Eco-mIL12, and the PBS control (Fig. 5D). TRP1^high^ CD8 T cells were specifically activated, as evidenced by an increase in CD25 expression post-virus injection (Fig. 5E). TRP1^high^ T cells exposed to A1-ZsG virus did not show a significant increase in CD25 expression compared to PBS controls, indicating that pMHC-targeted viruses can further fine-tune cell phenotype via delivery of a therapeutic cargo, as demonstrated here with tethered IL-12.

To explore changes in the tumor microenvironment that could explain observed differences in overall survival, tissues from a second cohort of mice were analyzed 6 days after treatment with virus, when tumor sizes began to diverge (Fig. 6, A and B). After A1-IL12 + aPD1 injection, tumor-infiltrating TRP1^high^ T cells significantly increased compared to injections of PBS + aPD1 and Eco-mIL12 (Fig. 6C). Additionally, significantly more IFNγ was detected in A1-IL12 + aPD1 tumor lysates, indicating that the tethered mIL-12 was initiating downstream signaling *in vivo* (Fig. 6D). One well-characterized impact of an IFNγ response in innate immune cells is the upregulation of MHC class II (MHC II) by macrophages (*49, 50*). We found that the ratio of tumor-infiltrating MHC II high to MHC II low macrophages was skewed towards an MHC II high response in mice injected with A1-mIL12 + aPD1 compared to PBS + aPD1 or A1-ZsG (Fig. 6E). Collectively, these results underscore that delivery of tethered mIL-12 via pMHC-targeted viruses impacts both adaptive and innate tumor-infiltrating immune cells, driving expansion of on-target cells and initiating a broader IFNγ -based immune response within the tumor microenvironment.

**Fig. 6.**
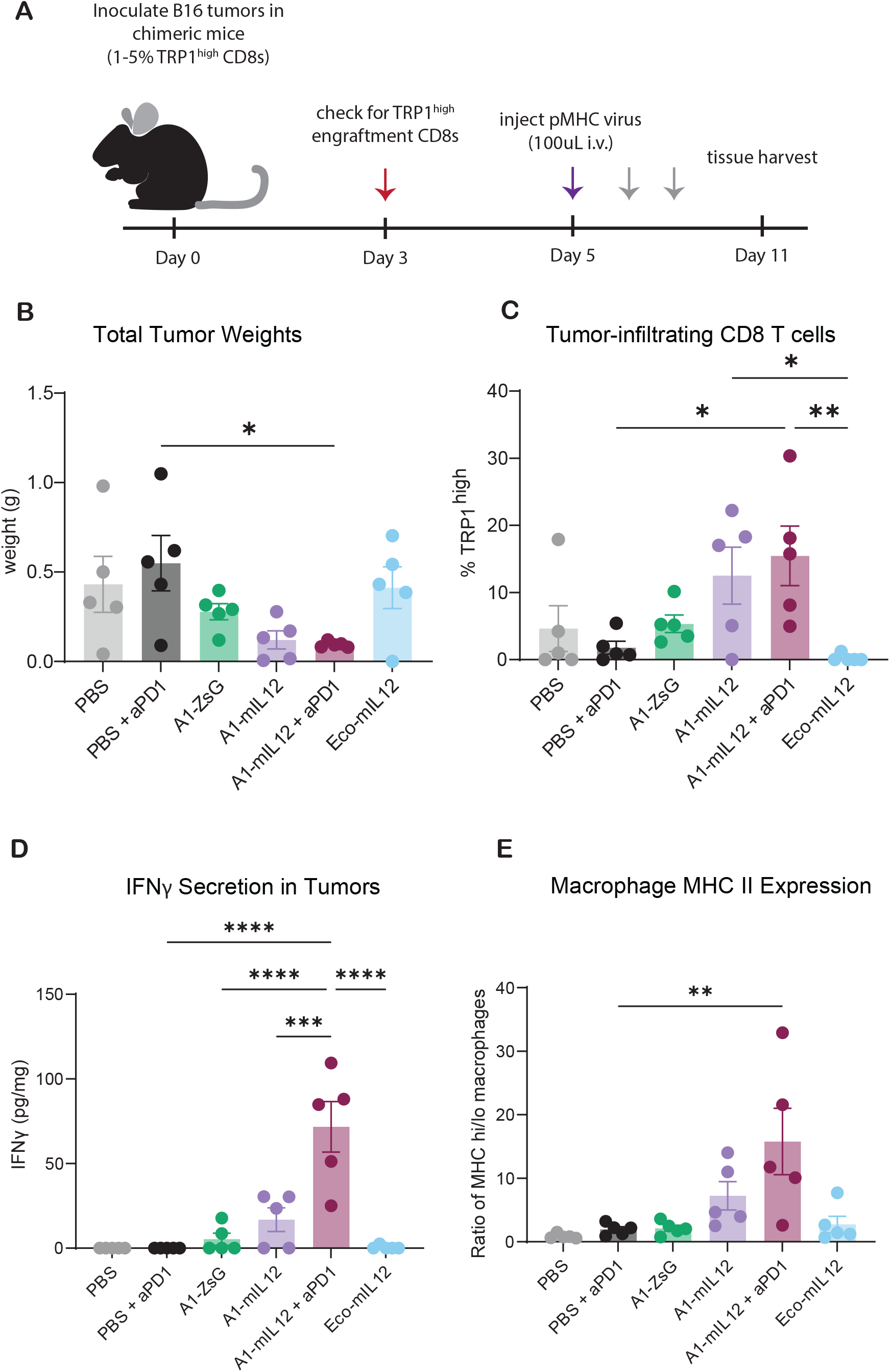
A1-mIL12 viruses recruit adaptive and innate immune cells in an anti-tumor response. (**A**) B6 mice were irradiated and partially reconstituted with TRP1^high^ lymphocytes. Two days later, 500,000 B16F10 tumor cells were inoculated subcutaneously. Three days later 100uL of targeted virus was injected i.v. via the tail vein. Grey arrows indicate i.p. injections of anti-PD-1 antibody (150ug) on days 7 and 10. Red arrows indicate a blood draw to check for TRP1^high^ T cell engraftment prior to virus injection. On Day 11, tissues were harvested for analysis via flow cytometry. (**B**) Tumor weights on Day 11. P values were calculated using Bonferroni-corrected one-way ANOVA. *p < 0.05 (**C**) Tumors were processed for flow cytometry and tumor-infiltrating TRP1^high^ T cells were quantified as a percentage of total tumor-infiltrating CD8s. Significance was calculated using a Bonferroni-corrected one-way ANOVA. *p < 0.05, ** p < 0.01 (**D**) IFNγ secretion was quantified in tumor lysates and normalized to total tumor weight. P values were determined using a Bonferroni-corrected one-way ANOVA. *** p < 0.001, **** p < 0.0001 (**E**) MHC II expression on tumor-associated macrophages was determined via flow cytometry. The ratio of MHC high to MHC low macrophages (MHC hi/lo) was calculated as a ratio of counts of each cell type. Significance was determined using a Bonferroni-corrected one-way ANOVA. ** p < 0.01

### CD8 T cells transduced *in vivo* with A1-mIL12 viruses can form long-term memory

One of the most promising aspects of engineered cell therapy is the potential to induce long-lasting memory with a single dose of treatment. To evaluate whether pMHC-pseudotyped viruses delivering mIL12 could generate immunologic memory, we increased the dose of viruses delivered systemically to B16F10 tumor-bearing mice partially reconstituted with TRP1^high^ T cells (Fig. 7A). Similar to previous *in vivo* transduction experiments, we saw specific expansion of the TRP1^high^ T cell population following addition of virus, with negligible transduction detected in off-target CD8s, CD4s, and non-T cell subpopulations (Fig. 7, B and C, and fig. S4, A and B). These results demonstrate that even at higher systemic doses, pMHC-targeted viruses remain targeted to antigen-specific T cells. At the same timepoint, we detected a significant increase in serum IFNγ when delivering mIL12 to tumor-specific T cells compared to non-specific delivery of IL-12 (Fig. 7D). Serum IL-12 levels were comparable between mice dosed with A1-mIL12 and Eco-mIL12 (Fig. 7E) suggesting that the IL-12 largely remains tethered to the cell surface. At this elevated dose of systemically delivered virus, mice receiving A1-mIL12 virus showed robust control of tumor growth, with 40% of mice alive at 40 days (Fig. 7F). Taken together, these data suggest that pMHC-targeted viruses are an efficient means of delivering potent immunomodulatory cargoes that can be localized to an antigen-specific T cell population of interest.

**Fig. 7.**
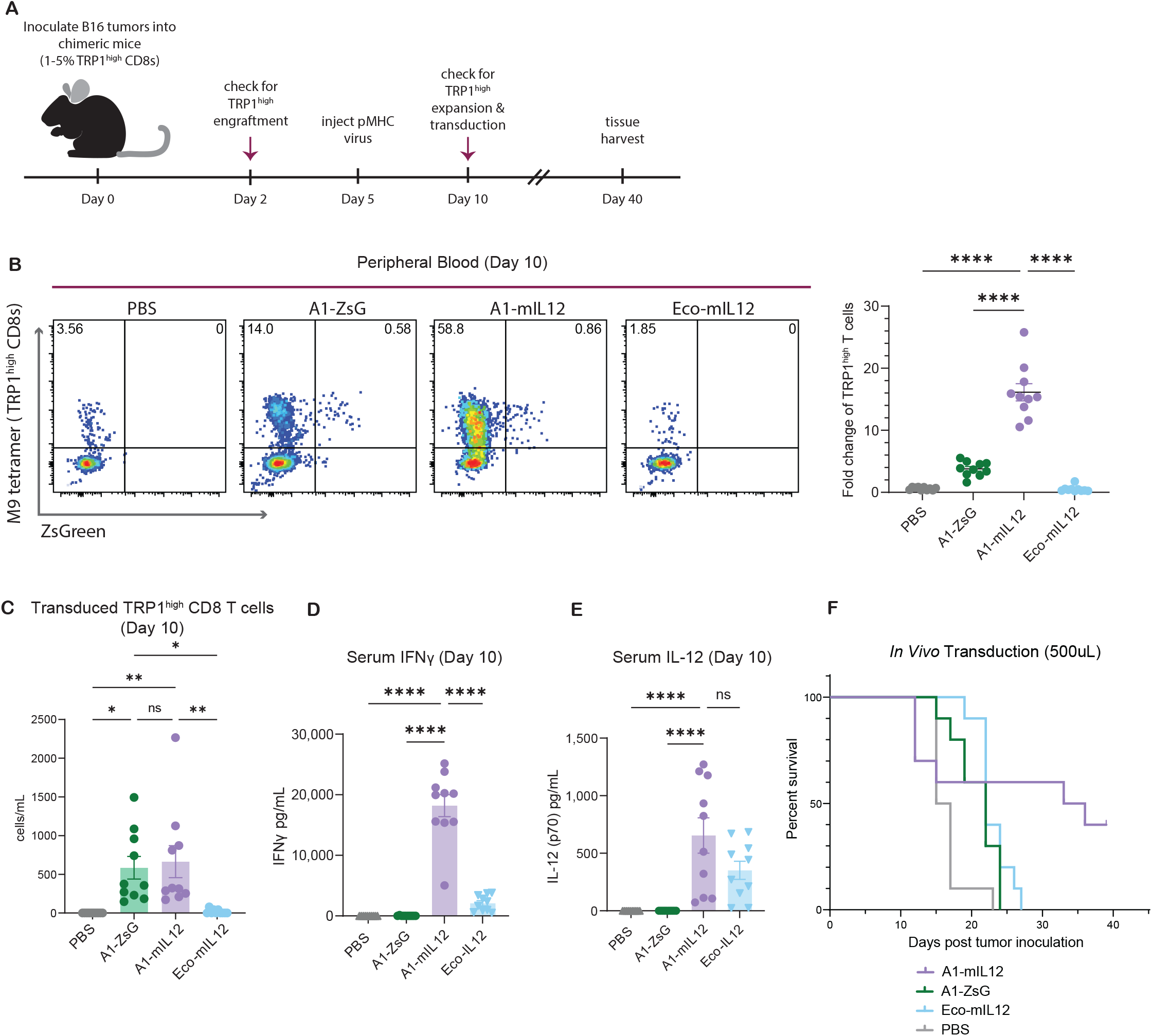
Increasing A1-mIL12 dose amplifies TRP1^high^ expansion and transduction *in vivo*. (**A**) 500,000 B16F10 tumor cells were inoculated subcutaneously on the left flank of mice reconstituted with TRP1^high^ lymphocytes. Three days later, a total of 500uL of virus was injected (300uL i.v. + 200uL i.p.). Maroon arrows indicate blood draws to check for the presence/expansion of TRP1^high^ T cells. At Day 40, surviving mice were sacrificed to determine whether transduced TRP1^high^ T cells were able to persist and traffic to tumors. (**B**) Gated on live CD8 T cells, from peripheral blood draw on Day 10. Representative plots from one mouse in each group. Annotated percentages are of total, live CD8 T cells. Fold change of TRP1^high^ T cells was calculated by comparing the frequency of TRP1^high^ T cells at Day 10 with their initial frequency at Day 2. P values were calculated using a Bonferroni-corrected one-way ANOVA. **** p < 0.0001 (**C**) Absolute counts of transduced TRP1^high^ T cells detected at Day 10 in peripheral blood. P values were determined using a Bonferroni-corrected one-way ANOVA. * p < 0.05, **p < 0.01 (**D**) On Day 10, IFNγ concentration in serum was determined by IFNγ ELISA. P values were assigned using a Bonferroni-corrected one-way ANOVA. **** p < 0.0001 (**E**) Samples from (D) were also assessed for presence of IL-12 in serum by IL-12 (p70) ELISA. P values were calculated using a Bonferroni-corrected one-way ANOVA. **** p < 0.0001 (**F**) Kaplan Meier curves for overall survival of mice in this study.

To determine whether transduced cells could contribute to immunologic memory, we evaluated the four remaining mice in the A1-mIL12 group at day 40 to assess the presence and persistence of transduced TRP1^high^ T cells in peripheral blood as well as in tumors (Fig. 8A). In two out of the four mice, the majority of CD8 T cells were TRP1^high^ T cells (Fig. 8B), and a detectable fraction of those TRP1^high^ T cells were stably transduced at Day 40 and exhibited an effector memory phenotype (CD44+CD62L-) (Fig. 8, C to E). These results emphasize that *in vivo* engineered, antigen-specific T cells can persist to become long-lived T cells following a single intravenous injection of pMHC-targeted virus. Together, these results underscore that pMHC-targeted viruses can be used to directly transduce antigen-specific T cells *in vivo* with a function-enhancing cargo, generating long-lived engineered cells which orchestrate a robust anti-tumor immune response.

**Fig. 8.**
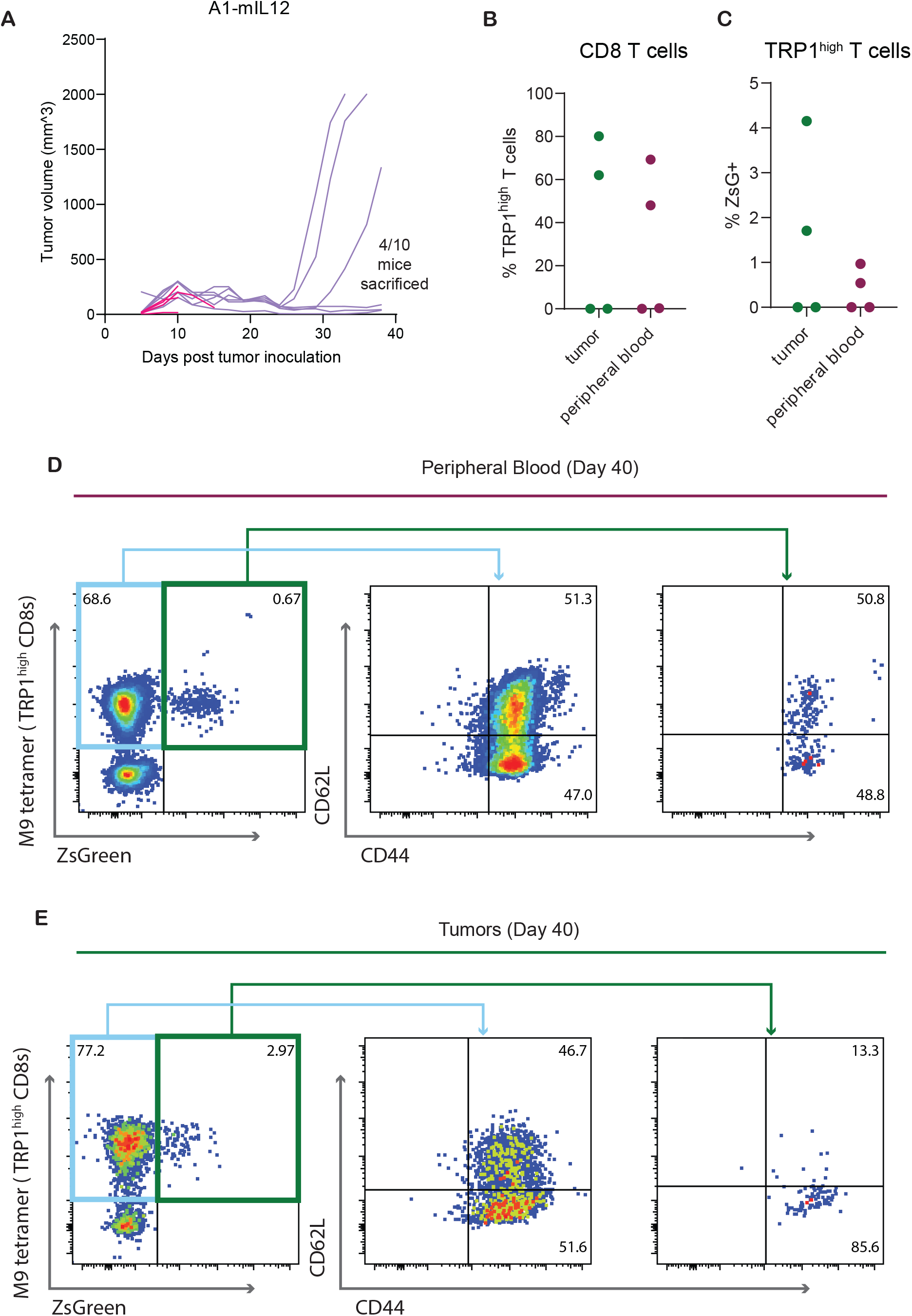
A single injection of A1-mIL12 virus is able to generate *in vivo* memory TRP1^high^ CD8 T cells in a subset of treated mice. (**A**) Tumor volume measurements from the A1-mIL12 treated group. Four of these mice were euthanized at early timepoints upon fulfillment of humane endpoint criteria due to initial treatment toxicities (shown in pink). Two mice were euthanized when tumor size reached maximum volume, and four mice maintained relatively small tumors until Day 40 when all remaining animals were sacrificed to determine persistence and trafficking of transduced TRP1^high^ T cells. (**B**) Frequencies of TRP1^high^ T cells in tumor and peripheral blood detected at Day 40. (**C**) Frequencies of transduced TRP1^high^ T cells in TRP1^high^ populations detected in (B). (**D-E**) Memory phenotypes of circulating (D) or tumor-infiltrating (E) TRP1^high^ T cells were characterized within untransduced or transduced cell populations. Naïve T cells = CD62L+CD44-, central memory T cells = CD62L+CD44+, effector memory T cells = CD62L-CD44+, stem cell-like memory = CD62L-CD44-. The first flow panel is gated on live CD8 T cells, with subsequent panels gated on either untransduced or transduced TRP1^high^ T cells as indicated by the boxes and arrows.

## DISCUSSION

The clinical success of TIL therapy and checkpoint blockade have underscored that pre-existing anti-tumor T cells present in patient T cell repertoires can mediate robust clinical responses. To fully leverage the potential of these endogenous, tumor-specific T cells, we demonstrate that pMHC-targeted retroviruses can be engineered to target antigen-specific CD8 T cells for *in vivo* delivery of function-enhancing genetic cargoes to modify T cell phenotype.

Our optimized pMHC pseudotyping strategy for gammaretroviruses facilitates efficient transduction, activation and expansion of primary anti-tumor CD8s. In TIL therapy, enriching for tumor-reactive T cells and minimizing bystander T cell involvement has been demonstrated to increase clinical efficacy (*51*). These data suggest that the ability of pMHC-targeted viruses to expand antigen-specific cells during the transduction process is a feature that could be important for the improved therapeutic outcomes we observe in both the *ex vivo* manufacturing and *in vivo* transduction models.

A major bottleneck in the development of cell therapy products is scaling the time-consuming and labor-intensive manufacturing process to meet growing clinical need. Because of these specialized infrastructure and skilled operator requirements, only a limited number of institutions are capable of manufacturing engineered cell therapies (*52*), further restricting patient access while increasing costs. Allogeneic cell products present one alternative to autologous cell engineering by providing off-the-shelf solutions. However, manufacturing allogeneic cell therapies still requires costly *ex vivo* manufacturing protocols similar to autologous cell therapies with the additional risk of generating alloreactive T cell responses. The enhanced specificity of pMHC-targeted viral vectors could bridge this gap by enabling *in vivo* engineering of autologous T cells. We validate that pMHC-targeted viruses injected systemically are able to specifically expand and transduce tumor-specific T cells at low initial frequencies *in vivo* with minimal off-target transduction.

In this work, we demonstrate delivery of an immunomodulatory cargo that catalyzes improved outcomes in tumor-bearing mice. We chose a tethered IL-12 construct as an ideal test cargo for our studies based on its narrow therapeutic index despite its ability to recruit multiple immune cell subtypes and orchestrate an anti-tumor response. Prior work demonstrates the efficacy but limiting toxicity of TILs engineered to secrete IL-12 (*28*). For this reason, efforts have been made to limit delivery of IL-12 to the tumor microenvironment (*48, 53*), by requiring a hypoxic microenvironment to induce IL-12 secretion (*54*), or by directly tethering the cytokine to endogenous as well as CAR T cells (*55*–*57*). We report that our antigen-specific virus could be an effective way to safely dose mIL12 systemically as only anti-tumor T cells are transduced with a spatially constrained construct. We expect that a wide variety of therapeutic proteins could be delivered using pMHC-targeted viruses, and future work could continue to explore whether certain cargoes are especially synergistic with our viral pseudotyping strategy that also activates and expands target T cells.

The modularity and generality of our vectors could be leveraged for the simultaneous delivery of cargoes to T cells recognizing a pool of antigens, further enhancing TIL therapy and reducing the potential for immune escape. Expression of single-chain trimers for viral targeting should be readily translated to all MHC alleles and can be incorporated into viral manufacturing without any additional steps beyond the inclusion of a gene encoding the single-chain pMHC. Tools to determine which pMHCs are recognized by tumor-specific TCRs already exist and could be leveraged to identify pMHCs that should be displayed as targeting molecules on the viral surface (*24, 58, 59*).

Although this demonstration of pMHC-targeted viruses was conducted within the setting of cancer immunotherapy, both the targeting and delivery components of our viral vectors are highly modular, making antigen-specific gene therapy using pMHC-targeted viral vectors a promising tool for development of cell therapies for a range of different diseases. Altogether, the pMHC-targeted vectors for gene delivery explored here could be powerful tools for improving the precision and quality of future cell therapy products.

## MATERIALS AND METHODS

### Study Design

The objective of this study was to evaluate pMHC-targeted viruses as gene delivery vectors in an immunocompetent model system. We first validated that pMHC-targeted viruses can specifically modify different antigen-specific CD8 T cell populations by generating pMHC-targeted gammaretroviruses and isolating transgenic CD8 T cells for *in vitro* transduction. Subsequently, we adoptively transferred these *ex vivo* engineered cells into mice bearing B16F10 tumors and monitored overall survival to determine if therapeutic efficacy could be achieved following antigen-specific delivery of a function-enhancing cargo to tumor-specific cells. Following an observed increase in overall survival for mice receiving A1-mIL12 transduced cells, we harvested cells for flow cytometry analysis to verify persistence of transduced cells. Finally, we demonstrate that pMHC-targeted retroviruses can be injected systemically for *in vivo* engineering of tumor-specific T cells present at a starting frequency of 1-5% of the total CD8 T cell repertoire. With this model, we measured overall survival, evaluated tumor-infiltrating cells via flow cytometry, and quantified cytokine secretion in tumors and serum.

### Plasmids

The plasmid pCL-Eco was a gift from Inder Verma (Addgene plasmid # 12371 ; http://n2t.net/addgene:12371 ; RRID:Addgene_12371) The plasmid pUMVC was a gift from Bob Weinberg (Addgene plasmid # 8449 ; http://n2t.net/addgene:8449 ; RRID:Addgene_8449). MSCV ZsGreen plasmid was cloned by inserting an IRES followed by the coding region of ZsGreen into the MSCV vector backbone, involving digestion with XhoI/EcoRI (NEB) and subsequent Gibson Assembly. pMD2.G was a gift from Didier Trono (Addgene plasmid # 12259 ; http://n2t.net/addgene:12259 ; RRID:Addgene_12259). The plasmid pMD2-VSVGmut was previously published and developed by our lab(*24*) (Addgene plasmid # 182229 ; http://n2t.net/addgene:182229 ; RRID:Addgene_182229). To generate MSCV IL-12, murine IL-12 was expressed as a tethered construct as previously described,(*48*) consisting of the coding sequence of a single chain murine IL-12 (a gift from Dane Wittrup) followed by a flexible G4S linker and residuces 247-268 of the transmembrane domain of murine CD80 (UniProt ID Q00609). This tethered mIL-12 construct was inserted into the MSCV ZsGreen plasmid directly upstream of IRES-ZsGreen via digestion of the vector with XhoI/EcoRI (NEB) followed by Gibson Assembly. Correct assembly was validated via Sanger and Oxford Nanopore sequencing.

### Generation of HEK producer cell lines

For each pMHC of interest, we generated HEK293T producer cell lines for stable mammalian cell expression of the targeting pMHC construct. To accomplish this, we cloned each single chain pMHC into the pHIV vector and generated VSVG pseudotyped lentiviruses delivering a pMHC of interest. Lentiviruses were produced by transfecting 80-95% confluent HEK293T cells with pHIV-pMHC, psPAX2, and pMD2.G (encoding VSVG) plasmids at a mass ratio of 5.6:3:1. Plasmids were complexed with TransIT-Lenti Transfection Reagent (Mirus Bio) at a DNA mass ratio of 3:1 in Opti-MEM (ThermoFisher). After a 10 minute incubation, DNA + TransIT-Lenti complexes were added dropwise to HEK293T cells. 48 hours later, HEK293T supernatant was collected and filtered using a 0.45 micron low protein-binding filter, and unconcentrated viral aliquots were stored at -80°C. These VSVG pseudotyped viruses and 8ug/mL of diethylaminoethyl-dextran (Sigma-Aldrich) were added to freshly split HEK293Ts. After 48hrs, transduction was assessed via flow cytometry by trypsonizing and staining the transduced HEK293T cells with antibodies specific for the pMHC of interest. A pure population of pMHChi HEK293T cells were then sorted using FACS (Sony M900). Following the sort, pMHChi HEK293T cells were expanded and frozen down to form producer cell line stocks.

### pMHC retrovirus production

pMHC-expressing HEK293T producer cell lines were thawed and passaged a minimum of 3 times in DMEM complete media, composed of DMEM (ATCC) supplemented with 10% fetal bovine serum and 100U/mL penicillin-streptomycin (Corning). Cells were then cultured to 80-95% confluency and transfected with pCL-Eco and an MSCV transfer plasmid (MSCV-ZsGreen or MSCV-mIL12) at a mass ratio of 3:5. Plasmids were complexed with TransIT-LT1 Transfection Reagent (Mirus Bio) at a mass ratio of 1:3 in OptiMEM (ThermoFisher). After incubating for 25 minutes, DNA + TransIT-LT1 complexes were added dropwise to HEK293T cells. Supernatants were collected at 48 hours post transfection and filtered with a 0.45um low protein-binding filter. Cell media was replaced with fresh DMEM complete media plus 25mM HEPES and a second collection was conducted at 72 hours post transfection.

Unconcentrated virus was immediately frozen at -80°C for future assays. Concentrated virus was produced by pooling supernatants from 48 and 72 hour collections and adding a PEG concentration cushion composed of 45% Poly(ethylene glycol) 6,000 in PBS at a volume of 1:3 PEG cushion to virus supernatant. Virus supernatant plus PEG cushion were stored at 4°C overnight and subsequently centrifuged at 1,500*g* for 45 minutes. The supernatant from this spin was discarded, and the PEG + virus pellet was resuspended in OptiMEM (ThermoFisher) at 1/100 of the volume of the harvest HEK supernatant. This concentrated virus was then aliquoted and stored at -80°C.

### Animals

C57BL/6J mice aged 6 to 8 weeks were purchased from Jackson Laboratories. All animal studies were performed in accordance with guidelines approved by the MIT Division of Comparative Medicine as well as the MIT Committee on Animal Care (Institutional Animal Care and Use Committee, protocol number 0621-032-24 and 2404000656) and in accordance with DFCI IACUC-approved protocols (14-019). TRP1^high^ TCR transgenic mice were bred in-house, and mice aged 8-24 weeks were used as T cell donors. 2C TCR and OT-I transgenic mice were gifts from the Spranger lab and bred in-house at the Koch Institute (MIT) mouse facility. Chimeric mice for *in vivo* pMHC-targeting were generated by irradiating C57BL/6J with 100 rads. 4-6 hours later, lymphocytes from TRP1^high^ mice were transferred intravenously via tail vein injection. All animal work was conducted in compliance with the NIH/NCI ethical guidelines for tumor-bearing animals.

### Isolation and transduction of murine CD8s

CD8s were isolated from spleens and inguinal lymph nodes of transgenic and B6 mice, with each strain processed separately. Tissues were pooled and filtered through a 70 micron filter to create single cell suspensions. Downstream processing was conducted following Easy Sep Mouse CD8+ T cell isolation kit (STEMCELL Technologies) or Miltenyi Biotec CD8a+ T Cell Isolation Kit for murine cells following manufacturer’s instructions (Miltenyi Biotec). Isolated CD8s were cultured at 1 million cells/mL in RPMI 1640 (ATCC) supplemented with 10% fetal bovine serum, 100U/mL penicillin-streptomycin (Corning), 50uM β-mercaptoethanol (ThermoFisher), 1% sodium pyruvate (ThermoFisher), 1% non-essential amino acids (ThermoFisher), and 10ng/mL murine IL-2 (Sigma-Aldrich). To transduce cells, 8ug/mL Polybrene (Santa Cruz Biotechnology) and concentrated virus were added followed by spinfection (1000*g* at 32°C for 1.5hrs).

### Tumor inoculations for *in vivo* experiments

B16F10 cells were cultured until 80-90% confluent, then trypsonized, washed with PBS, and resuspended in Hank’s balanced salt solution (HBSS) at 2 x 10^6^ cells/mL. Mice were shaved one day prior to inoculations and 250uL (5 x 10^6^) cells were injected subcutaneously into the left flank. Before injections, mice were randomly assigned to treatment groups. Baseline weight was measured on injection day for injections of volumes less than or equal to 100uL or one day after injections of larger volume. Body weight was monitored 2-3 times every week. In survival experiments, tumor size was measured every 2-4 days. Mice were euthanized when tumor volume exceeded 2000mm^3^, tumors showed evidence of ulceration, or weight decreased 20% relative to baseline.

### *Ex vivo* transduction for adoptive transfer experiments

CD8 T cells to be transduced by Eco-targeted gammaretroviruses were isolated three days prior to adoptive transfer and were activated with 10uL/M cells using Dynabeads Mouse T-activator CD3/CD28 beads (ThermoFisher) for 24 hours. Two days before adoptive transfer, all cells were transduced following the above protocol. Equivalent numbers of pre-activated or freshly isolated T cells were transduced across all groups. Virus doses were standardized across groups so that the same number of particles/M cells was added in every condition, where number of particles was quantified based on p30 ng/mL determined by MuLV Core Antigen ELISA (Cell Biolabs). Across all three replicate experiments, a dose of 7.9 x 10^3^ – 1.2 x 10^4^ ng p30/M cells was used to transduce each group. For cells transduced by A1-targeted viruses, no pre-activation was required, and 5ug/mL InVivoMAb anti-CD28 antibody (BioXCell) was added in conjunction with concentrated virus prior to spinfection. On transfer day, cells were counted and stained to assess viability, activation, and percent transduction. All cells in the resultant cell product for each group were transferred, with a minimum of 800,000 transduced cells injected per mouse and not exceeding a maximum dose of 10.3M total cells/mouse.

### *In vivo* transduction experiments

B16F10 tumors were inoculated in chimeric C57BL/6J mice as described above. Two to three days after tumor inoculation, baseline frequency of TRP1^high^ CD8 T cells in peripheral blood was quantified via a retro-orbital bleed. Five days after tumor inoculation, 100-300uL of virus was injected i.v. into the tail vein with an additional 200uL injected i.p. in studies where 500uL total virus was delivered. Five days after virus injection, peripheral blood was again assayed to determine expansion, activation, and transduction of TRP1^high^ CD8 T cells.

In experiments where a combination of virus plus checkpoint blockade was tested, aPD1 (clone RMP1-14, BioXCell) was administered at a 150ug dose in endotoxin-free PBS delivered i.p. at the timepoints specified for a maximum of four total doses.

### Tissue collection for histology

Livers and spleens were collected and placed in fixative, Z-fix (Fisher Sci, NC9378601), for 30 to 45 minutes then left in 30% sucrose in PBS overnight at 4°C. Tissues were embedded in OCT blocks and frozen by floating on liquid nitrogen before storage in -80°C. Sections were acquired using a Leica CM3050S Cryostat sectioned at -20°C at 7 μm per slide. Staining was performed by washing with 0.1% Triton-X 100 in PBS for 3 x 15 minutes then staining for CD3-PE (1:100, Biolegend, 100205) overnight at room temp. Slides were washed in 0.1% Triton-X 100 in PBS for 3 x 15 minutes then stained with DAPI (1:2000) for 30 minutes at room temperature in the dark. Slides were washed once in PBS before being mounted in Prolong Gold Antifade Mountant (ThermoFisher, P36930) and imaged using an Olympus IX73 fluorescent microscope using a Hamamatsu Orca-Spark C11440-36U camera.

### Antibodies and flow cytometry

For flow cytometry analysis, cells were first washed with PBS then stained 1:500 with LIVE/DEAD Fixable Viability Dye (ThermoFisher) and 1:200 with purified anti-mouse CD16/32 (Biolegend, clone 93) for 15 minutes on ice. If CD8 T cells were isolated, no purified anti-mouse CD16/32 was used. Following an additional wash step, this time in FACS buffer (PBS mixed with 0.5% BSA and 2mM EDTA), samples were stained for 15 minutes on ice with fluorescently labeled antibodies at 1:200 dilution from the stock solution. Cells were then washed once with FACS buffer prior to analysis on a Cytoflex S (Beckman) or Sony Biotechnology SP6800 Spectral Analyzer. For experiments where absolute cell counts were calculated, CountBright™ Absolute Counting Beads (ThermoFisher) were added to each well just prior to analysis. M9 tetramers, displaying the TRP1 epitope TAPDNLGYM in the context of MHC H2-D^b^, were produced in-house: single-chain MHC monomers were expressed in High Five cells (ThermoFisher), purified, and biotinylated. Monomers were then mixed with PE streptavidin (Biolegend) at a ratio of 5:1 in PBS and incubated on ice for 10 minutes. When relevant, tetramers were added to cells concurrently with antibodies which were then collectively incubated for 30 minutes at room temperature.

Tumors were processed for flow cytometry by removing the entire tumor, using scissors to manually dissociate the tissue. Samples were then incubated in RPMI supplemented with tumor digestion enzymes (Miltenyi tumor dissociation kit, Cat. No. 130-096-730) at 37°C for 30 mins. Following digestion, samples were filtered through a 40um strainer to ensure generation of single-cell suspensions. Spleens were processed by filtering through a 40um strainer and performing one round of ACK lysis (ThermoFisher). Lymph nodes were ground through 40um filters and resuspended for downstream staining. Peripheral blood was subjected to two rounds of ACK lysis prior to subsequent staining.

### pSTAT4 intracellular staining

CD8 T cells from a TRP1^high^ mouse were isolated as described above and transduced with A1-ZsG or A1-mIL12 viruses following the same *in vitro* transduction protocol outlined previously. Two days after transduction, cells were fixed by adding an equal volume of pre-warmed BD Phosflow Fix Buffer I (BD Biosciences) to the media and incubating for 10 minutes at 37°C. Cells were washed twice with FACS buffer and fixed with BD Phosflow Perm Buffer III for 30 minutes on ice. After permeabilization, cells were washed twice with FACS buffer and stained with 647 anti-pSTAT4 (pY693) (1:50) for 1 hour on ice in the dark. Following staining, cells were washed with FACS buffer and acquired on a Cytoflex S (Beckman).

### IFNγ and IL-12 ELISAs

ELISA MAX™ Deluxe Set Mouse IFN-γ (Biolegend) and ELISA MAX™ Standard Set Mouse IL-12/IL-23 (p40) (Biolegend) were used to quantify IFNγ production and untethered IL-12 respectively. Assays were conducted according to the manufacturer’s instructions.

For ELISAs of cell supernatants from *in vitro* cultures, supernatants were collected 48 hours after transduction and frozen at -80 until the time of assay.

Tissue samples for IFNγ analysis were collected 11 days post tumor inoculation and flash frozen on the day of tissue harvest. To make tumor lysate, tumors were thawed in 100-300uL RIPA lysis buffer with a protease inhibitor (Millipore Sigma) and phosphatase inhibitor cocktail (Cell Signaling Technology) and transferred to bead beating tubes (OMNI). Tubes were placed in a Bead Ruptor and tumors were mechanically dissociated for 30 seconds. Tumor lysate was then centrifuged at 13,000*g* for 20 minutes at 4C. Supernatant was collected and frozen at -80°C for subsequent processing. Pierce BCA Protein Assay kits (ThermoFisher) were used to determine tumor lysate protein concentration for normalization.

### Software

Graphs were generated using GraphPad Prism 10. Flow cytometry data were analyzed by FlowJo (10.10.0). Figures were composed and compiled using Adobe Illustrator.

### Statistical analysis

GraphPad Prism 10 was used to conduct statistical analyses. Results were considered significant if p < 0.05. * p < 0.05; ** p < 0.01; *** p < 0.001; **** p < 0.0001

## Supplementary Materials

Figs. S1 to S4

## Acknowledgments

For their technical support, we thank the Koch Institute’s Robert A. Swanson (1969) Biotechnology Center, particularly the Flow Cytometry Facility. We thank Beth E. Grace, Vidit Bhandarkar, and Stefani Spranger for their reagents and expertise in developing pMHC-targeted viruses for the 2C and OT-1 model systems. We also thank Anisha Datta for her assistance in developing ELISA analysis pipelines as well as Nishant Singh for his protocol to make pMHC tetramers.

## Funding

This research was supported in part by the Koch Institute Frontier Research Program, the Damon Runyon-Rachleff Innovation Award (58S-20), the Gates Foundation, and NIH DP2 (DP2-AI158126) to M.E.B.. E.J.K.X was supported by a fellowship from the Ludwig Center at MIT’s Koch Institute and through Singapore MIT Alliance for Research and Technology (SMART): Critical Analytics for Manufacturing Personalised-Medicine (CAMP) Inter-Disciplinary Research Group. B.E.S. (T32GM007753) and A.G. (T32GM144273) were supported by the National Institute of General Medical Sciences. S.K.D. was supported by a Technology Impact Award from the Cancer Research Institute, the Ludwig Center at Harvard, Break Through Cancer, NIH R01AI158488, R01AI169188, and is a Member of the Parker Institute for Cancer Immunotherapy.

## Author contributions

Conceptualization: E.J.K.X, S.K.D., and M.E.B

Methodology: E.J.K.X, M.J.W, M.T.H., L.Q.

Investigation: E.J.K.X., B.E.S., W.C.A., M.J.W., B.L., M.T.H., L.Q., J.D., A.G., Q.H.Z., and C.R.P.

Analysis and visualization: E.J.K.X., M.T.H., and L.Q.

Funding acquisition: M.D., S.K.D, and M.E.B

Supervision: M.D., S.K.D, and M.E.B

Writing (original draft): E.J.K.X., M.D., S.K.D., and M.E.B.

Writing (review and editing): all authors

## Competing interests

M.E.B. is a founder, consultant, and equity holder of Kelonia Therapeutics and Abata Therapeutics. S.K.D. received research funding unrelated to this project from Novartis, Bristol-Myers Squibb, Takeda, and is a founder, science advisory board member and equity holder in Kojin and has equity in Axxis Bio. M.D. has research funding from Eli Lilly; he has received consulting fees from Genentech, ORIC Pharmaceuticals, Partner Therapeutics, SQZ Biotech, AzurRx, Eli Lilly, Mallinckrodt Pharmaceuticals, Aditum, Foghorn Therapeutics, Palleon, and Moderna; and he is a member of the Scientific Advisory Board for Neoleukin Therapeutics, Veravas and Cerberus Therapeutics and has equity in Axxis Bio. C.S.D. is currently employed by and is an equity holder of Kelonia Therapeutics. The remaining authors declare no competing interests.

## Data and materials availability

All data generated in this study are included in the main text or supplementary figures. Plasmids generated for this investigation are available upon request. Any additional information required to reproduce this work can be made available upon request by the corresponding authors, M.E.B and S.K.D.

