## Supplemental Figures for "Peptide-MHC-targeted retroviruses enable *in vivo* expansion and gene delivery to tumor-specific T cells"

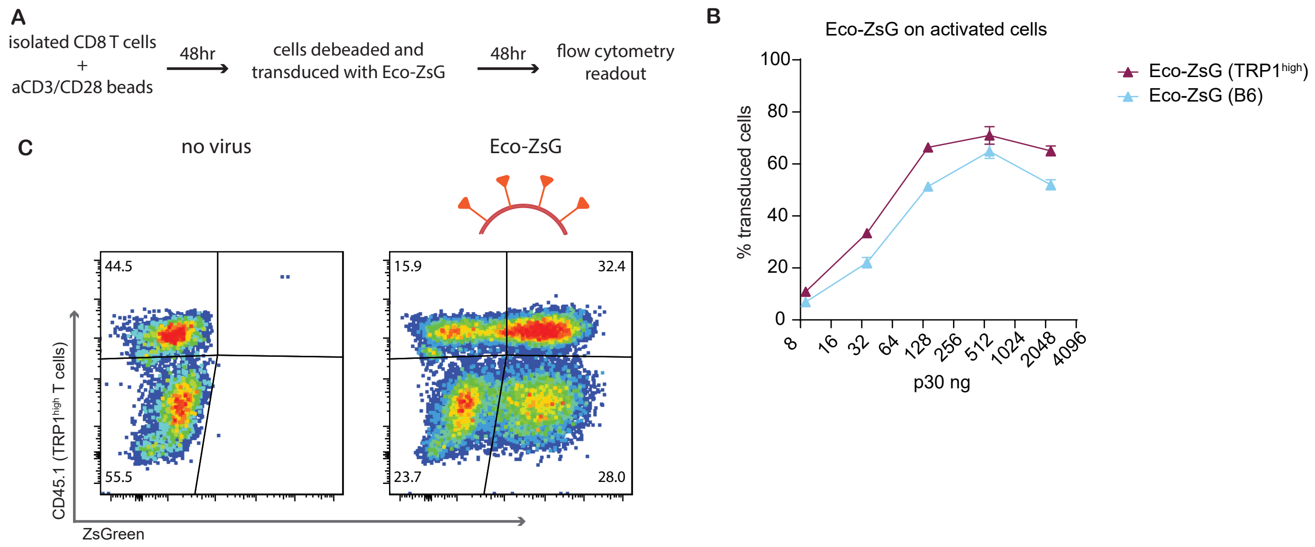


**Fig. S1. Eco-ZsGreen viruses efficiently transduce activated CD8 T cells in an antigen-independent manner.** (A) Design of experiment evaluating potential of Eco-ZsG to transduce activated cells. Isolated TRP1high and B6 CD8 T cells were mixed at a 1:1 ratio and activated with a-CD3/CD28 beads. After two days, cells were transduced with virus, and results were analyzed via flow 48 hours later. (B) Expression of ZsGreen as a percentage of total TRP1high or B6 cells. Representative of n = 3 technical replicates. (C) Representative plots from data summarized in (B) displaying the transduction observed in the no virus condition (left) or addition of Eco-ZsG (right).

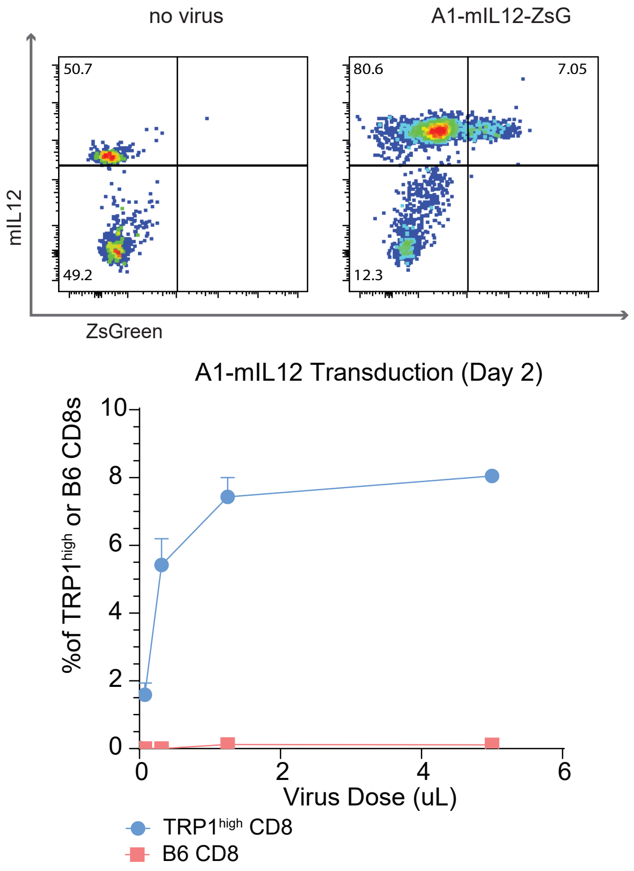


**Fig. S2. A1-targeted viruses retain their ability to expand and transduce TRP1high T cells specifically when delivering a function-enhancing cargo, mIL12.** (A) Representative flow plots of transduction achieved after adding A1-mIL12 virus (right) to a 1:1 mixture of TRP1high and B6 CD8 T cells measured after 48 hours. Single experiment from n = 3 biological replicates. (B) Summary plot of (A) where n = 3 technical replicates.

A

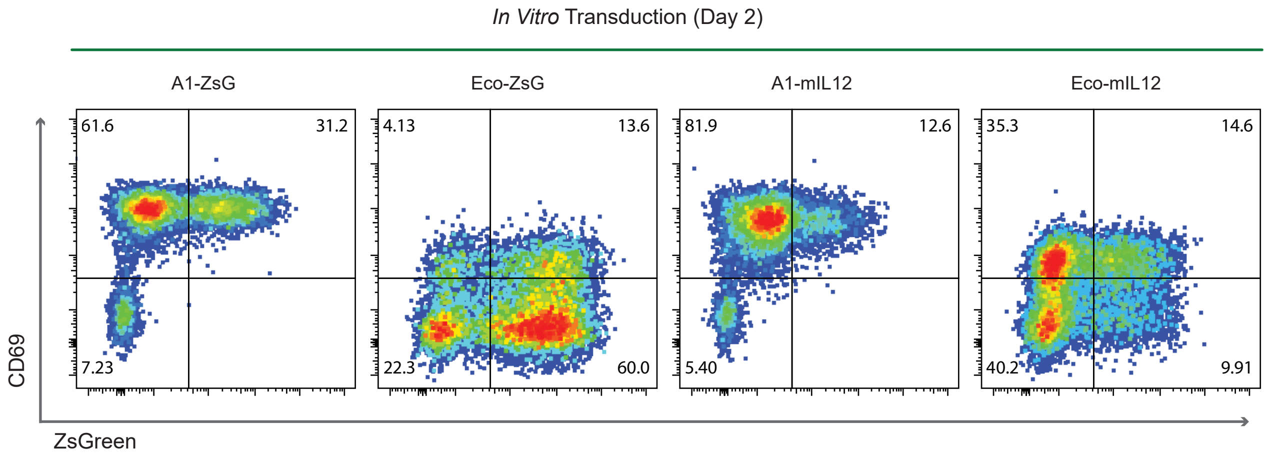


B

| Virus | Cells | Total cells transferred/mouse | Transduced cells transferred/mouse |
|---|---|---|---|
| A1-ZsG | TRP1[high] | 10.3M | 2.9M |
| Eco-ZsG | TRP1[high] (pre-activated) | 10.3M | 7.2M |
| A1-mIL12 | TRP1[high] | 10.3M | 0.9M |
| Eco-mIL12 | B6 (pre-activated) | 10.3M | 2.1M |

**Fig. S3. TRP1high T cells are transduced and activated during ex vivo production process prior to adoptive transfer.** (A) Evaluation of transduction, as a measure of ZsGreen expression, and activation, as a measure of CD69 expression, of each group of in vitro transduced cells two days after addition of virus, preceding adoptive transfer. (B) Table outlining the cells (TRP1high or B6, pre-activated or freshly isolated) that were transduced in each group. A total of 10.3M cells were transferred per mouse in each treatment group, but due to inherent differences in the number of transducing units per mL of each virus, different quantities of total transduced cells per mouse were transferred.

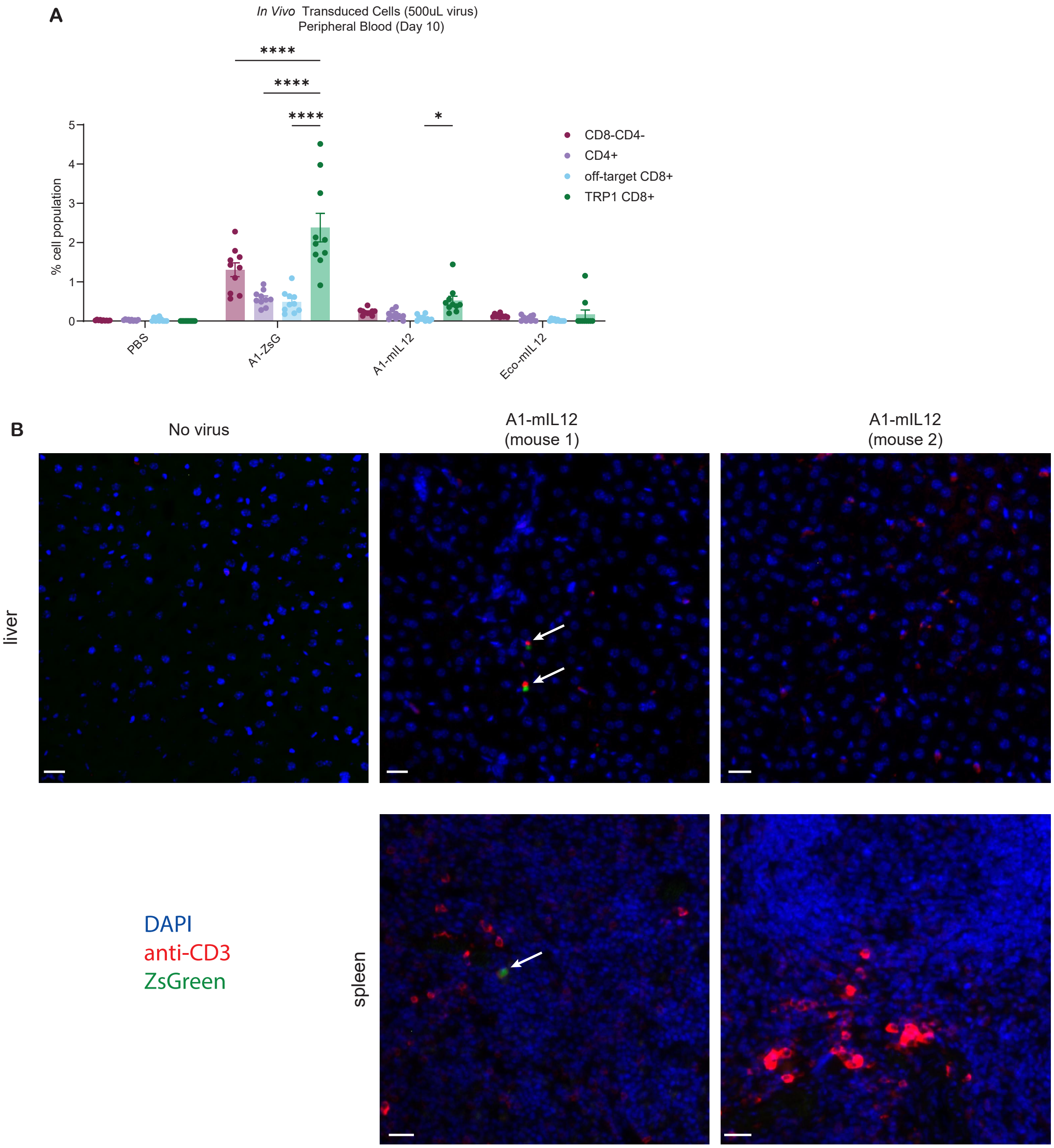


**Fig. S4. Minimal off-target transduction is detected even at high, systemic doses of A1-mIL12 virus.** (A) ZsGreen+ cells detected in peripheral blood at Day 10, 5 days after injection of 500uL of virus. Transduction is quantified as a frequency of the total cell population, defined by the indicated markers. P values were determined by Bonferroni-corrected two-way ANOVA. *$p < 0.05$, ****$p < 0.0001$ (B) Tissues from mice in the same experiment as (A) were harvested at Day 40, fixed, and sectioned for subsequent analysis. Shown are representative images from an untreated mouse and two mice treated with A1-mIL12, where each image is from a different mouse. Tissues were stained with DAPI (blue) and anti-CD3 (red) alongside ZsGreen (green). Images shown are from livers (top row) and spleens (bottom row). White arrows highlight instances of ZsGreen. Selected images were chosen to maximize display of detectable ZsGreen in the tissue of interest. Scale bars shown in white are 20um.
